# *mimic*INT: a workflow for microbe-host protein interaction inference

**DOI:** 10.1101/2022.11.04.515250

**Authors:** Sébastien A. Choteau, Kevin Maldonado, Aurélie Bergon, Marceau Cristianini, Mégane Boujeant, Lilian Drets, Christine Brun, Lionel Spinelli, Andreas Zanzoni

## Abstract

**Background:** The increasing incidence of emerging infectious diseases is posing serious global threats. Therefore, there is a clear need for developing computational methods that can assist and speed up experimental research to better characterize the molecular mechanisms of microbial infections.

**Methods:** In this context, we developed *mimic*INT, an open-source computational workflow for large-scale protein-protein interaction inference between microbe and human by detecting putative molecular mimicry elements mediating the interaction with host proteins: short linear motifs (SLiMs) and host-like globular domains. *mimic*INT exploits these putative elements to infer the interaction with human proteins by using known templates of domain-domain and SLiM-domain interaction templates. *mimic*INT also provides *(i)* robust Monte-Carlo simulations to assess the statistical significance of SLiM detection which suffers from false positives, and *(ii)* an interaction specificity filter to account for differences between motif-binding domains of the same family. We have also made *mimic*INT available via a web server.

**Results:** In two use cases, *mimic*INT can identify potential interfaces in experimentally detected interaction between pathogenic *Escherichia coli* type-3 secreted effectors and human proteins and infer biologically relevant interactions between Marburg virus and human proteins.

**Conclusions:** The *mimic*INT workflow can be instrumental to better understand the molecular details of microbe-host interactions.

## Introduction

Most pathogens interact with their hosts to reach an advantageous niche and ensure their successful dissemination. For instance, viruses interfere with important host-cell processes through protein-protein interactions to coordinate their life cycle (1). It has been shown that host cell networks subversion by pathogen proteins can be achieved through interface mimicry of endogenous interactions (i.e., interaction between host proteins) (2,3). This strategy relies on the presence in pathogen protein sequences of host-like elements, such as globular domains and short linear motifs (SLiMs), that can mediate the interaction with host proteins (4–6).

Over the last years, many computational methods have been developed to predict pathogen-host protein interactions, some of which are based on the detection of sequence or structural mimicry elements (7–9). Such approaches allowed, for instance, to suggest potential molecular mechanisms underlying the implication of gastrointestinal bacteria in human cancer (10,11) or to discriminate between viral strains with different oncogenic potentials (12), thus showing that protein-protein interaction predictions can be instrumental in untangling microbe-host disease associations. Nevertheless, the source code of many of these tools is not freely available to the community (e.g., (11–13)) providing the predictions through a database (e.g., (12)), or can be only used through a web interface (14,15), thus limiting reproducibility and tool usability.

In this context, and inspired by our previous work (10), we have developed the *mimic*INT workflow, and its webserver companion *mimic*INTweb (https://mimicintweb.tagc.univ-amu.fr), to enable large-scale interaction inference between microbe and human proteins based on the detection of host-like elements and the use of experimentally identified interaction templates (16,17).

## Methods

### Implementation

*mimic*INT detects putative molecular mimicry elements in microbe sequences of interest that can mediate the interaction with host proteins (Figure 1). *mimic*INT is written in Python and R languages and exploits the Snakemake workflow manager for automated execution (18). It consists of four main steps: *(i)* the detection of host-like elements in microbe sequences; *(ii)* the collection of domains on the host protein *(iii)*; the interaction inferences between microbe and host proteins; and *(iv)* the functional enrichment analysis on the list of inferred host interactors.

**Figure 1.**
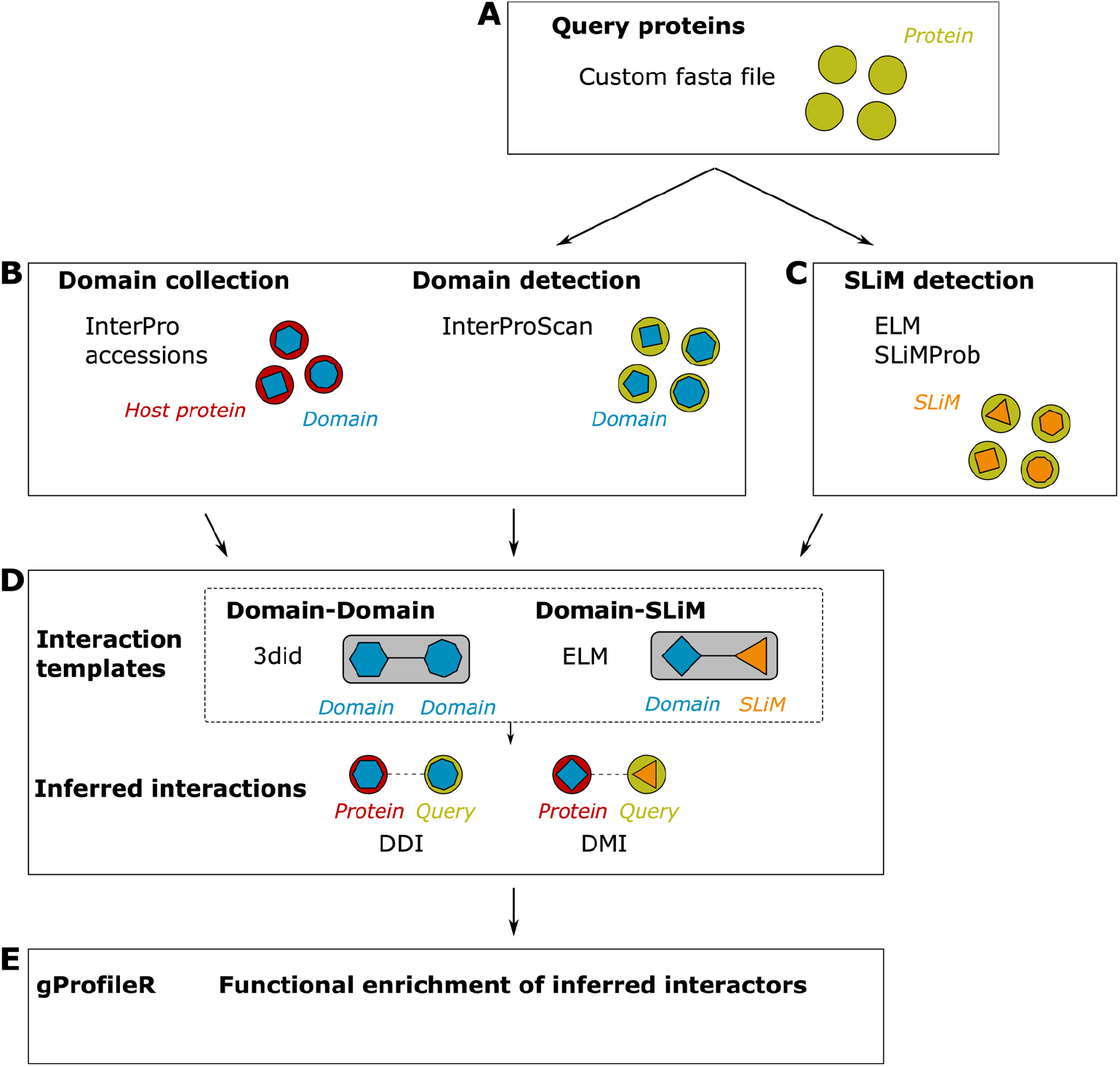
Overview of the *mimic*INT workflow. By providing a fasta file of protein sequences of the query species (e.g., microbe sequences) (A), *mimic*INT allows identifying both the domain- (B) and SLiM- (C) mediated interfaces of interactions. Using publicly available templates of interactions, *mimic*INT infers the interactions between the proteins of the query and target (i.e., host) species (D). Finally, it provides a list of functional annotations significantly enriched in inferred protein targets (E).

In the first step, *mimic*INT takes the FASTA-formatted sequences of microbe proteins (*e*.*g*., viral or other pathogen proteins susceptible to be found at the pathogen-host interface) as input to detect host-like elements: domains and SLiMs. The domain identification is performed by the InterProScan stand-alone version (19) using the domain signatures from the InterPro database (20). By default, *mimic*INT retains InterProScan matches with an *E*-value below 10^−5^, a common threshold value used for detecting profile-based domain signatures in protein sequences in the context of interaction inference (21). The host-like SLiM detection exploits the motif definitions available in the ELM database (17) and is carried out by the SLiMProb tool from the SLiMSuite software package (22). As SLiMs are usually located in disordered regions (23), SLiMProb uses the IUPred algorithm (24) to compute the disorder propensity of each amino acid in the query sequences and generates an average disorder propensity score for every detected SLiM occurrence. For SLiM detection, the default IUPred disorder propensity threshold is set to 0.2, a value commonly used to limit false negatives (22,25), and the minimum size of the predicted disorder region is set to 5, which is the optimal size to detect true positive SLiM occurrences (26). Nevertheless, the user can choose all running parameters for the host-like element detection in the *mimic*INT configuration file.

In the second step, *mimic*INT gathers the domain annotations of the host proteins from the InterPro database through a REST API query.

In the third step, *mimic*INT infers the interactions between host and microbe proteins. This analysis takes as input the list of known interaction templates collected from two resources: *(i)* the 3did database (16), a collection of domain-domain interactions extracted from three-dimensional protein structures (27), and *(ii)* the ELM database (17) that provides a list of experimentally identified SLIM-domain interactions in Eukaryotes. The inference procedure checks whether any of the microbe proteins contain at least one domain or SLIM for which an interaction template is available. In this case, it infers the interaction between the given protein and all the host proteins containing the cognate domain (*i*.*e*., the interacting domain in the template). As motif-binding domains of the same group, like SH3 or PDZ, show different interaction specificities (28), we have implemented a previously proposed strategy (29) to take these differences into account (see below the sub-section “Computation of the motif-binding domain similarity scores”). This approach assigns a “domain score” that can be used to rank, or filter inferred SLiM-domain interactions. Once this step is completed, the inferred interactions are stored in both tab-delimited and JSON files to facilitate the import in other applications, such as Cytoscape (30).

In the final step, to identify the host cellular functions potentially targeted by the pathogen proteins, *mimic*INT executes a functional enrichment analysis of host-inferred interactors. This analysis statistically assesses the over-representation of functional categories, such as Gene Ontology terms and biological pathways (e.g., KEGG and Reactome), using the g:Profiler R client (31).

Given the degenerate nature of SLiMs (23), their detection is prone to generate false positive occurrences. For this reason, we implemented an optional sub-workflow that, using Monte-Carlo simulations, assesses the probability of a given SLiM to occur by chance in query sequences and, thus, can be used to filter out potential false positives (5) (see below the sub-section “Statistical significance of the SLiMs detected on the microbe sequences”).

To ease deployment and ensure reproducibility and scalability on high-performance computing infrastructures, *mimic*INT is provided as a containerized application based on Docker and Singularity (32,33).

### Computation of the motif-binding domain similarity scores

To identify motif-binding domains that can be specifically associated with a given ELM motif class, we use the same strategy proposed by Weatheritt et al. in 2012 (29), which assumes that a domain significantly similar to a known motif-binding domain should also bind the same motif. We first compiled a list of experimentally identified motif-binding domains from the original list from Weatheritt et al. complemented by more recent annotations from the ELM database (17) (August 2020). Obsolete ELM class identifiers from Weatheritt et al. were mapped to current ELM identifiers using the “Renamed ELM classes” file (http://elm.eu.org/infos/browse_renamed.tsv) and duplicated domain annotations were removed. In total, we collected 538 domains in 415 human proteins known to bind 212 ELM motif classes (73% of the 290 motif classes present in ELM, August 2020). The sequences of these 415 annotated proteins were fetched from UniprotKB (34). We next fetched the sequences of 1452 reference Eukaryota proteomes (22,262,113 protein sequences in total) from UniprotKB (August 2020). We removed redundancy using the CD-HIT algorithm (35) to generate a database of 21,414,544 non-identical sequences. We used the GOPHER tool (36) from the SLiMSuite package (22) to identify orthologous sequences of the annotated proteins in the database of non-identical eukaryotic sequences by reciprocal BLAST best hits. Selected orthologous proteins were aligned using the multiple sequence alignment algorithm Clustal Omega (v. 1.2.4) (37). Once the position of the motif-binding domain was identified within the alignment, we removed aligned domains with indels covering >10% of the annotated domain sequence. We iteratively realigned the sequences until a set of proteins was identified with <10% indels coverage. In total, we selected 701 multiple sequence alignments used as input for generating domain-specific HMM profiles with the *hmmbuild* program from the HMMER package v.3.1.1 (38). Subsequently, we scanned a representative set of the human proteome (20,350 “reviewed” sequences from UniprotKB) with the domain-specific HMMs using the *hmmsearch* program. We used an *E*-value cutoff of 0.01 to select the best hits and we rejected those hits covering less than 90% of the annotated motif-binding domain sequence length. Finally, the *E*-value of the best-scoring domain was converted into a domain similarity score using the iELM script downloaded from http://elmint.embl.de/program_file/ (29). Doing so, we computed at least one motif-binding domain similarity score for 1,461 human proteins.

### Statistical significance of the SLiMs detected on the microbe sequences

To assess the probability of a given motif to occur by chance in microbe sequences, we implemented a previously proposed approach (5) to randomly shuffle the disordered regions of each sequence of a microbe of interest to generate a large set of randomized microbe proteins. The number of shuffled sequences to be generated by *mimic*INT can be chosen by the user in the corresponding configuration file (see the *mimic*INT online documentation for more details). By default, *mimic*INT creates two sets of 100,000 randomly shuffled proteins (one set for each IUPred disorder propensity prediction mode, i.e. *short* and *long*), with the assumption that the input sequences belong to the same microbe species or strain. Once the shuffled sequences are generated, the occurrences of each detected motif are compared in each microbe input sequence to the occurrences observed in the corresponding set of shuffled sequences. To compute the probability (*P*) of each detected motif to occur by chance, *mimic*INT counts the number of times (*m*) out of the shuffled sequences (*N*) where there is at least the same number of instances of the given motif in the input sequence:

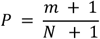

For example, if a given motif occurs twice in the input sequence, the methods count how many times the same motif is detected at least twice in the corresponding set of randomly shuffled sequences. The lower the value of *P*, the rarer the instances, thus suggesting that the given motif can be likely functional. In this work, we set the significant threshold equal 0.1, as reported in (5).

### Webserver

The *mimic*INTweb server allows users not familiar with the command-line interface to run the *mimic*INT workflow through an easy-to-use web interface. The number of input sequences is limited to 50. A step-by-step tutorial is available on the *mimic*INTweb site (https://mimicintweb.tagc.univ-amu.fr/tutorial). The *mimic*INTweb server uses the Django framework (version 2.2.1 under Python 3.12.4) as web app core to manage URL routing, HTML rendering, authentication, administration, and backend logic. The Django component has been complemented with two additional application layers to guarantee server performances and security: Gunicorn (version 22.0.0) as web server gateway interface, and Nginx (version 1.25) as reverse proxy server.

### Operation

The *mimic*INT workflow can be run on a Linux-based computer with at least 32 GB RAM and it has been successfully used on Ubuntu (16.04 and higher) and CentOS (7.4) distributions. The following software is required: Python (3.6 or higher), Snakemake (6.5 or higher), Docker (18.09 or higher) and IUPred (version 1.0). The workflow can be also deployed on high-performance computing (HPC) clusters. In this case, the Singularity application (2.5 or higher) is required. More detailed information can be found on the *mimicI*NT GitHub repository (https://github.com/TAGC-NetworkBiology/mimicINT). The *mimic*INTweb server can be accessed from Linux, Windows or Mac OS based systems, and it has been tested with the following browsers: Chrome, Firefox and Safari.

### Results

We sought to evaluate the ability of *mimic*INT to correctly infer SLiM-domain interactions, as this inference can generate many false positives (29), using the default parameters for SLiM detection (see Implementation). To do so, we used as controls two datasets of established motif-mediated interactions (MDI) from the ELM database (17): *(i)* 103 interactions between 87 viral and 44 human proteins (vMDI); *(ii)* 31 interactions between 16 bacterial and 23 human proteins (bMDI). We were able to correctly infer most of these interactions (91 vMDI, true positive rate = 88.3%; 21 bMDI, true positive rate = 67.7%). Notably, almost all the correctly inferred interactions have a domain score above 0.4 (87 out of the 91 vMDI, 19 out of 21 bMDI). As the availability of negative SLiM-mediated interaction datasets is very limited (17,29,39), we estimated the false positive rate (FPR) by applying *mimic*INT to two sets of randomly generated interaction sets (degree-controlled, vMDI_*rnd*_ and bMDI_*rnd*_, respectively). Thirty-four vMDI_*rnd*_ and 7 bMDI_*rnd*_ were inferred as motif-mediated (FPR = 33% and FPR = 23%, respectively). We next annotated the human proteins in the two random sets with domain scores. We kept only interactions for which the domain score was above 0.4 (29), thereby reducing the number of random interactions predicted as motif-mediated to 9 (FPR = 8.7%) for vMDI_*rnd*_ and 2 (FPR = 6.4%) for bMDI_*rnd*_. Finally, we tested *mimic*INT on two sets of experimentally verified negative protein interactions from the Negatome 2.0 database (40): 37 viral-human and 4 bacterial-human interactions. Only two virus-human negative interactions (5.4%) were inferred as motif-mediated by *mimic*INT.

In light of these results, we used *mimic*INT in two tasks: (i) the identification of putative interfaces in experimentally identified interactions between secreted effectors from the enteropathogenic *Escherichia coli serotype O157:H7* (EHEC) and human proteins; (ii) the inference of interactions between human and the Marburg virus (MARV) proteins, an emerging infectious agent for which experimental protein interaction data is scarce.

### Interface identification in the EHEC-human protein interaction network

We collected 83 interactions between 24 EHEC secreted effectors and 74 human proteins by querying (January 2022) the IMEx consortium databases (41) via the PSICQUIC interface (42). We gathered the sequences of EHEC effectors from UniprotKB in FASTA format and ran *mimic*INT with default parameters. We computed the motif probabilities using the dedicated sub-workflow by performing 100,000 randomizations. We were able to identify a putative interaction interface for 26 of the 83 experimental EHEC-human interactions (31.3%) (Figure 2A), which is higher than the number of interactions with identified putative interfaces in a degree-controlled randomized network. Most of the putative interfaces were identified using motif-domain interaction templates (MDI), namely 24 interactions, whereas the putative interfaces for 9 interactions were identified with domain-domain interaction templates (DDI). Interestingly, we identified putative interfaces with both MDI and DDI templates for 7 interactions (Figure 2A). Among the interactions with MDI interfaces, almost all have a motif probability below 0.1 (23 interactions, see Supplementary File 1). Seven interactions have a domain score above 0.4 (29.2%) and their cognate motifs show a motif probability lower than 0.1 (Figure 2B). This suggests that most of the identified putative interfaces can be considered as high confidence. To further support these inferences, we sought to verify whether the 26 putative interfaces corresponded to experimentally identified binding regions. To do so, we collected the biological features (i.e. “binding-associated region”, “necessary binding region”, “sufficient binding region”)(43) reported in the interaction records downloaded via the PSICQUIC interface, and we found that for half of the interactions with an inferred interface (13 interactions, 11 MDI and 2 DDI) there is supporting experimental evidence for at least one of the interaction partners (Figure 2C). For 7 interactions (27%), *mimic*INT inferred correctly the interface elements of both EHEC effectors and human proteins. For the other 4 interactions, the experimental evidence supports the EHEC effector interface element only (see Supplementary File 1). Importantly, the 11 MDI inferences can be considered of high confidence as they have either a motif probability < 0.1 or a domain score > 0.4. Overall, these results indicate that high confidence *mimic*INT inferred interaction can identify *bona fide* interaction interfaces.

**Figure 2.**
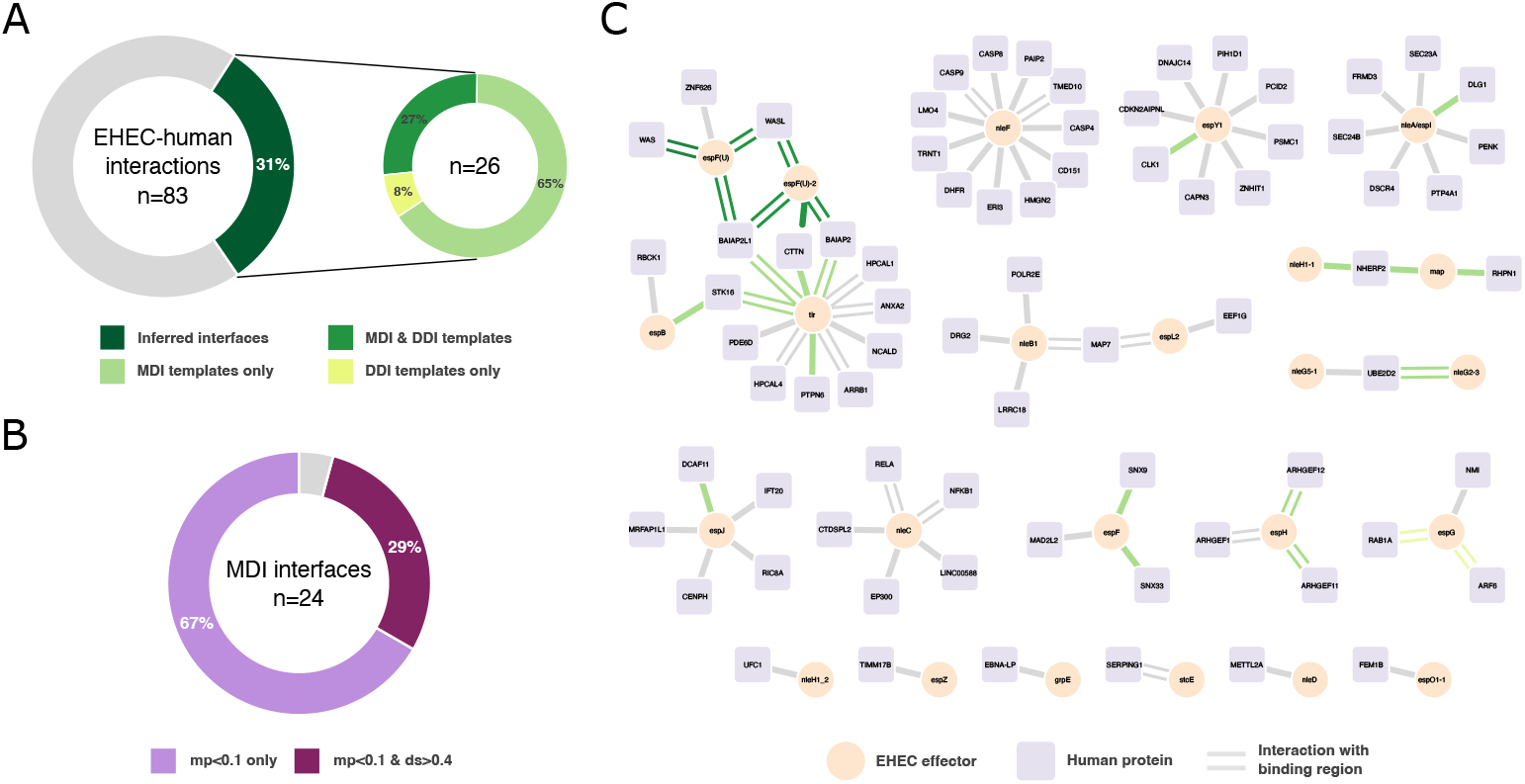
Application of the *mimic*INT workflow to identify potential interaction interfaces. **(A)** Proportion of experimentally determined interaction between EHEC secreted effectors and human proteins with at least one putative interface inferred by mimicINT (left). Split proportion of EHEC-human interactions with a putative interface according to the interaction templates: motif-domain (MDI) and domain-domain (DDI). **(B)** Proportion of EHEC-human interactions with high-confidence MDI-inferred interfaces based on the computed motif probability (mp) and domain score (ds). See main text for more details. (C) Network representations of the interactions between EHEC secreted effectors (circle nodes) and human proteins (square nodes). Edges represented as parallel lines indicate interactions with experimentally identified binding regions in at least one of the interaction partners. Coloured edges represent interactions with at least one putative inferred interface by *mimic*INT. The network was generated using Cytoscape (30).

### MARV-human protein interaction inference

We downloaded MARV protein sequences (7 proteins, Proteome ID: UP000180448, January 2022) from UniprotKB in FASTA format and ran *mimic*INT with default parameters. We also computed the motif probabilities using the dedicated sub-workflow by performing 100,000 randomizations.

In total, we inferred 11,431 interactions between 7 MARV and 2757 human proteins (see Supplementary File 2). Most of the inferred interactions, namely 10,101, are motif-domain interactions (MDI) between 7 MARV and 2324 human proteins, and the remaining 1,339 are domain-domain interactions (DDI) between 5 MARV and 479 human proteins (9 interactions were inferred with both MDI and DDI templates). The functional enrichment analysis performed by *mimic*INT on the full list of inferred host interactors returned 975 enriched annotations at FDR<0.01 (see Supplementary File 2). We further filtered out the functional categories annotating less than 5 or more than 500 proteins, obtaining a list of 763 enriched annotations (241 GO biological processes, 63 GO Cellular components, 6 CORUM complexes, 130 KEGG and 237 Reactome pathways, see Supplementary File 2), which points towards cellular processes and pathways related to viral infection and immune system (Table 1). By applying the default thresholds on motif probabilities and domain scores on inferred MDI, we defined a set of 535 high-confidence MDI interactions between 7 MARV and 419 human proteins. We combined this set with the inferred interaction using DDI templates and ran a functional enrichment analysis on a list of 891 human interactors returning 908 enriched annotations at FDR<0.01. As above, after filtering on the size of functional categories, we obtained 743 enriched annotations (287 GO biological processes, 57 GO Cellular components, 1 CORUM complexes, 141 KEGG and 257 Reactome pathways, see Supplementary File 2). Interestingly, 27% of the enriched GO biological process annotations (77 out of 287) are related to infection and immunity (44), and notably 8 out of the 10 most enriched.

**Table 1.**
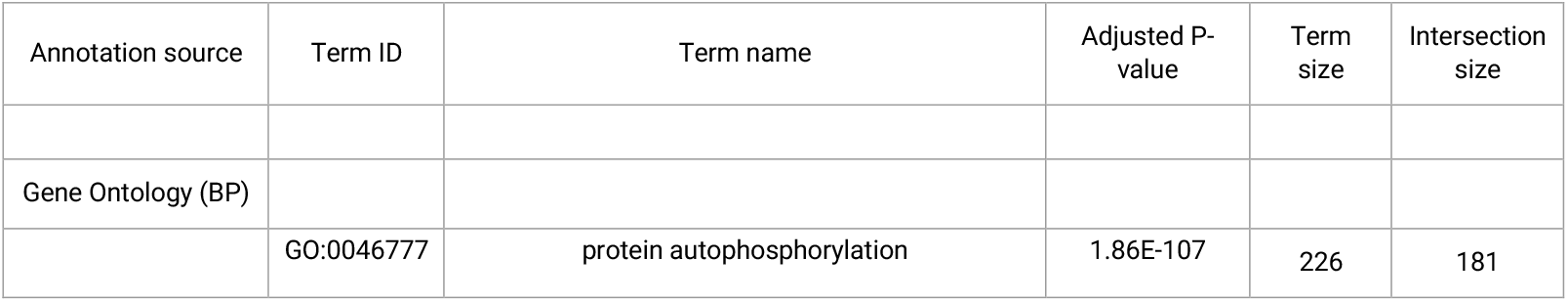
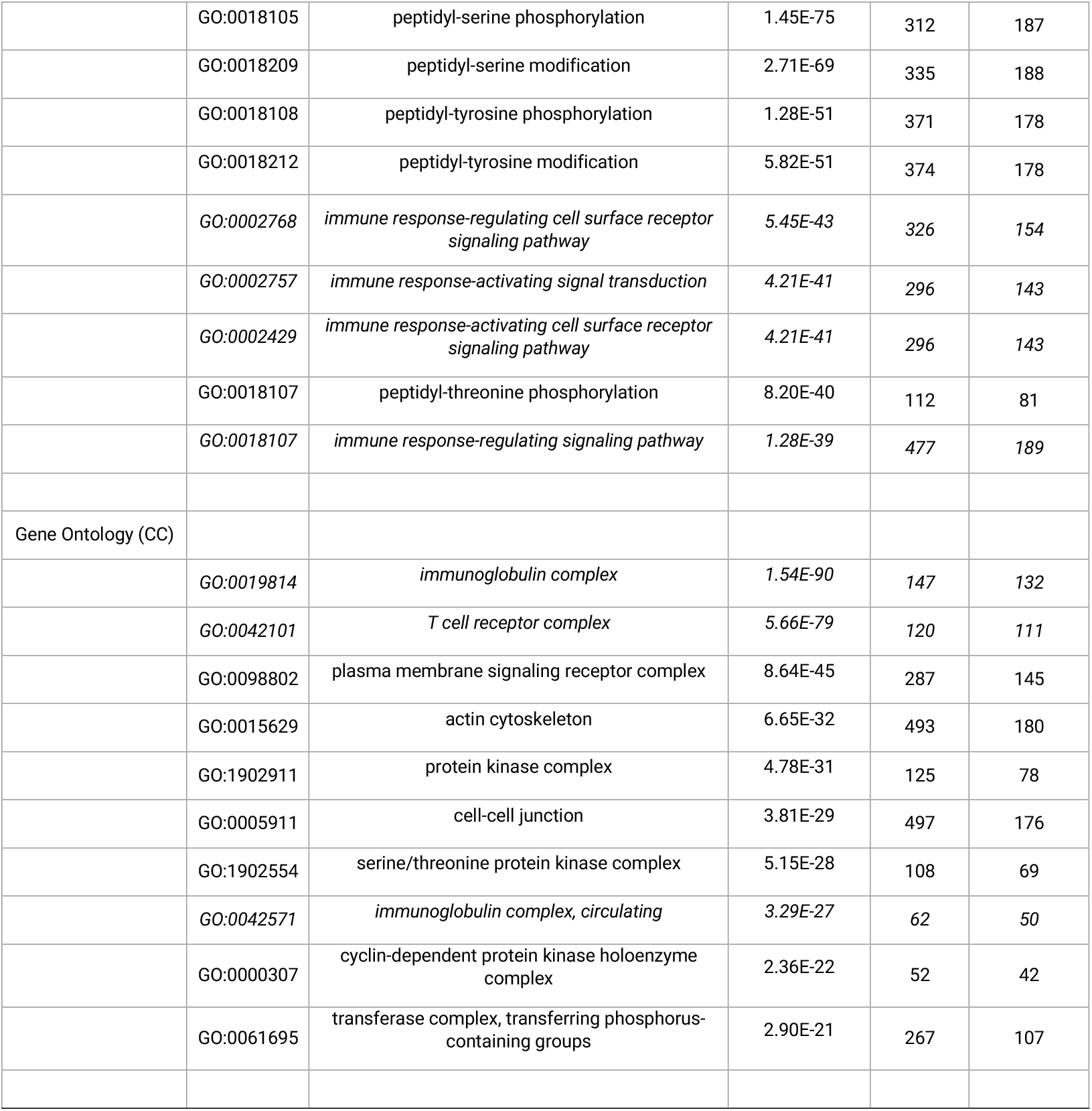
Summary results of the functional enrichment analysis performed by *mimic*INT on the 2685 human proteins inferred as interactors of MARV proteins. The top 10 most enriched terms are shown for Gene Ontology Biological Process (BP) and Cellular Component (CC) terms. For each enriched term the following information is reported: term identifier, term name, adjusted P-value, number of human proteins annotated with the given term in the statistical background (term size), number of inferred interactors annotated with the given term (intersection size). The terms reported in italic are related to viruses, infection and immunity according to Garcia-Moreno and colleagues (44).

| Annotation source | Term ID | Term name | Adjusted P-value | Term size | Intersection size |
| --- | --- | --- | --- | --- | --- |
| Gene Ontology (BP) |  |  |  |  |  |
|  | GO:0046777 | protein autophosphorylation | 1.86E-107 | 226 | 181 |
|  | GO:0018105 | peptidyl-serine phosphorylation | 1.45E-75 | 312 | 187 |
|  | GO:0018209 | peptidyl-serine modification | 2.71E-69 | 335 | 188 |
|  | GO:0018108 | peptidyl-tyrosine phosphorylation | 1.28E-51 | 371 | 178 |
|  | GO:0018212 | peptidyl-tyrosine modification | 5.82E-51 | 374 | 178 |
|  | *GO:0002768* | *immune response-regulating cell surface receptor signaling pathway* | *5.45E-43* | *326* | *154* |
|  | *GO:0002757* | *immune response-activating signal transduction* | *4.21E-41* | *296* | *143* |
|  | *GO:0002429* | *immune response-activating cell surface receptor signaling pathway* | *4.21E-41* | *296* | *143* |
|  | GO:0018107 | peptidyl-threonine phosphorylation | 8.20E-40 | 112 | 81 |
|  | *GO:0018107* | *immune response-regulating signaling pathway* | *1.28E-39* | *477* | *189* |
| Gene Ontology (CC) |  |  |  |  |  |
|  | *GO:0019814* | *immunoglobulin complex* | *1.54E-90* | *147* | *132* |
|  | *GO:0042101* | *T cell receptor complex* | *5.66E-79* | *120* | *111* |
|  | GO:0098802 | plasma membrane signaling receptor complex | 8.64E-45 | 287 | 145 |
|  | GO:0015629 | actin cytoskeleton | 6.65E-32 | 493 | 180 |
|  | GO:1902911 | protein kinase complex | 4.78E-31 | 125 | 78 |
|  | GO:0005911 | cell-cell junction | 3.81E-29 | 497 | 176 |
|  | GO:1902554 | serine/threonine protein kinase complex | 5.15E-28 | 108 | 69 |
|  | *GO:0042571* | *immunoglobulin complex, circulating* | *3.29E-27* | *62* | *50* |
|  | GO:0000307 | cyclin-dependent protein kinase holoenzyme complex | 2.36E-22 | 52 | 42 |
|  | GO:0061695 | transferase complex, transferring phosphorus-containing groups | 2.90E-21 | 267 | 107 |

Overall, these results reinforce the biological relevance of the inferred interactions, particularly those considered of higher confidence.

## Discussion

We have developed *mimic*INT, an open-source computational workflow enabling large-scale interaction inference between microbe and host proteins. In the first use case presented here, we show that *mimic*INT can identify *bona fide* interaction interfaces in an experimentally generated interaction network between secreted pathogenic bacterial effectors and human proteins. Notably, we also successfully used it to identify interaction interfaces between commensal bacterial effectors and human proteins in a large-scale interaction dataset generated by yeast two-hybrid (45). In the second use case, we used *mimic*INT to infer the interactions between viral and human proteins which are biologically relevant given the results of the functional enrichment analysis.

Although we developed *mimic*INT as a tool to infer protein interactions between microbe and human proteins, it can be used on any organisms whose proteins bear either domains or motifs with known interaction templates (e.g., human, mouse or fruit fly). For instance, we have recently used *mimic*INT to generate the first interactome of small human peptides encoded by short Open Reading Frames (sORFs)(46). Nevertheless, the only limitation of the workflow is the availability of motif-domain and domain-domain templates, which depends on the curation efforts done by teams maintaining the corresponding source database (i.e., ELM and 3did).

Finally, compared to other similar tools (47), *mimic*INT provides two functionalities to define high-confidence inferred interactions based on motif-domain templates, that is the computation of (i) motif probabilities and of (ii) motif-binding domain similarity scores. As shown in the use cases, the application of these two strategies supports the identification of bona fide interaction interfaces in the EHEC-human interaction network and the biological relevance of the inferred MARV-human interactions.

All in all, given the increasing frequency of (re-)emerging infectious diseases and the accumulating evidence on the fundamental role played by microbes in chronic diseases (48,49), there is no doubt that *mimic*INT will be useful to better understand the molecular details of the microbe-host relationships.

## Data availability

Supplementary Data are available from: https://doi.org/10.5281/zenodo.14614802

Data are available under the terms of the <u>Creative Commons Attribution 4.0 International license</u> (CC-BY 4.0).

## Software availability

Workflow source code available from: https://github.com/TAGC-NetworkBiology/mimicINT

Archived workflow source code at time of publication: https://doi.org/10.5281/zenodo.12078119

Webserver available from: https://mimicintweb.tagc.univ-amu.fr/

Webserver source code available from: https://github.com/TAGC-NetworkBiology/mimicINTweb

Archived webserver source code at time of publication: https://doi.org/10.5281/zenodo.14548031

License: <u>GNU General Public Licence, V3</u>.

## Acknowledgements

The authors thank Paul de Boissier for helping in the early development of the workflow and Fabrice Lopez for technical advice. The authors are also grateful to the members of the DIME project for fruitful scientific discussions. Centre de Calcul Intensif d’Aix-Marseille is acknowledged for granting access to its high-performance computing resources.

*Author contributions*: Conceptualization: S.A.C, C.B., L.S. and A.Z. Methodology: S.A.C., L.S. and A.Z. Software: S.A.C., K.M., A.B., M.C., L.D., L.S. and A.Z. Formal Analysis: S.A.C. and A.Z. Investigation: S.A.C., M.B. and A.Z. Writing – original draft: S.A.C. and A.Z. Writing – review & editing: C.B., L.S. and A.Z. Visualization: S.A.C. and A.Z. Supervision: C.B., L.S. and A.Z. Project Administration: C.B., L.S. and A.Z. Funding Acquisition: C.B and A.Z.

